# Limited cross strain CMV protection by a bivalent (gB/pp65) vaccine is based on an impaired gB antibody response and evasion of pp65 cell mediated immunity

**DOI:** 10.64898/2026.10.28.762556

**Authors:** K. Yeon Choi, Yushu Qin, Nadia El-Hamdi, Alistair McGregor

## Abstract

Congenital CMV (cCMV) is a leading cause of hearing loss and cognitive impairment in newborns and a vaccine is a high priority. The guinea pig with species-specific guinea pig cytomegalovirus (GPCMV) is the only small animal model for cCMV. Prior GPCMV vaccine studies have focused on prototype strain 22122 and not evaluated cross-strain protection, an important vaccine benchmark. The essential viral entry gB glycoprotein is an immunodominant antibody target and considered a corner stone of a CMV vaccine. Trimeric gB antigen induces potent neutralizing antibody titers but a clinical strain of GPCMV (TAMYC) is highly cell-associated limiting gB antibody cross-strain protection. In HCMV and GPCMV tegument protein pp65 is a major T cell antigen and important for convalescent immunity. A defective adenovirus (Ad) vaccine strategy encoding GPCMV gB and pp65 (AdgB+AdGP83) was evaluated for cross-strain protection. Vaccinated animals induced a cell-mediated response via IFNγ ELISPOT assay against pp65 peptides conserved between strains. Additionally, the combined vaccine produced similar anti-gB and prefusion gB titers to historical sera from a standalone trimeric gB vaccine (AdgB). However, antibody response to gB(AD-1) oligomerization domain was impaired in the combined vaccine compared to historical AdgB vaccine sera which resulted in reduced neutralizing titer that were more similar to naturally convalescent GPCMV infected animals. In a pathogenicity study, vaccinated animals had partial protection against clinical TAMYC GPCMV strain challenge (SQ) compared to non-vaccinated control group. However, the vaccine failed to prevent viremia or virus dissemination to target organs, including the salivary glands, a notable limitation. Potentially, TAMYC virus evades T cell response against pp65 by dramatically reducing levels of pp65 expression compared to prototype 22122 strain. Overall, results indicate that a combination vaccine offers limited cross-strain protection despite prior success of separate pp65 or gB vaccines against GPCMV (22122 strain) and conservation of antigens between strains.

## 3. Introduction

Human cytomegalovirus (HCMV) is a ubiquitous betaherpesvirus that establishes a life-long persistent infection with the ability to cross the placenta and cause congenital disease in utero [1]. Congenital cytomegalovirus (cCMV) in surviving newborns results in vision and cognitive impairment as well as sensorineural hearing loss (SNHL) and a potential risk factor for autism [2,3]. In USA and Europe, cCMV rate ranges from approximately 0.5-1.2% in newborn babies with up to 30% of hearing loss in children attributed to cCMV [4–6]. In regions of high CMV seroprevalence, the rate of cCMV increases up to 6.1% in some developing countries [7]. Consequently, a vaccine is considered a high priority but is complicated by the existence of multiple viral strains [2]. Convalescent HCMV natural immunity from a primary infection is insufficient to prevent re-infection by a new viral strain or risk of cCMV [8]. The immunological correlates for complete protection against cCMV infection are poorly defined and preclinical animal model studies enable important insights into CMV disease and development of intervention or therapeutic strategies [9]. HCMV species-specificity complicates studies in animal models, which require the use of species-specific animal CMV. The guinea pig is the only small animal model for cCMV with guinea pig cytomegalovirus (GPCMV) causing congenital disease in newborn pups similar to human and various strategies against CMV have been evaluated in this model [10,11].

HCMV viral glycoprotein complexes on the virion that are required for direct cell entry (gB, gM/gN, and gH/gL/gO) are antibody targets [12]. An additional gH/gL based pentamer complex (PC) in association with gB enables cell entry by an endocytic pathway [13,14]. Guinea pig CMV (GPCMV) encodes functional homolog glycoprotein complexes (gB, gH/gL/gO, gM/gN) essential for GPCMV infection and these complexes are neutralizing target antigens [10,15–17]. GPCMV also encodes a homolog PC necessary for endocytic pathway of infection in conjunction with gB [16]. Importantly, GPCMV uses similar cell receptors to HCMV for direct (PDGFRA) and endocytic (NRP2) pathways of cell entry [18]. In both HCMV and GPCMV, the gB glycoprotein is an immunodominant neutralizing antibody target and essential for infection of all cell types as the fusogenic viral glycoprotein [15,16,19–24]. However, a gB (HCMV) subunit vaccine, despite evoking high antibody titers, exhibited only approximately 50% efficacy in clinical trials [25,26]. GPCMV gB subunit vaccine strategies in the guinea pig model fail to fully protect against cCMV and lack cross-strain protection [27] exhibiting efficacy similar to that in human gB vaccine trials [10]. In both HCMV and GPCMV, gB antibodies are less effective at neutralizing virus infection of non-fibroblast cells (epithelial and endothelial cells) where cell entry is via the endocytic pathway [23,28–30]. In GPCMV, a trimeric capable gB vaccine (AdgB) resulted in better neutralizing titers on both fibroblast and non-fibroblast cells and improved efficacy against GPCMV prototype strain 22122 in a non-pregnant and cCMV protection models [23,24]. Prefusion gB (prefgB) has recently been characterized for HCMV and potentially an immune response to this novel antigen could improve the antibody-based response against gB [24,31]. However, antibody response to gB (AD-1) oligomerization domain is possibly considered a key neutralizing target antigen response [32].

The majority of GPCMV vaccine studies have focused on protection against prototype strain 22122 which was extensively passaged on fibroblast cells (>100) upon original isolation in the 1950s [33]. In HCMV, passage of clinical strains on fibroblasts resulted in numerous modifications to the genome compared to low pass clinical strains [34–36]. Additionally, despite the ability of epithelial cell or salivary gland stock derived virus to cause cCMV, a focus only on 22122 strain does not accomplish an important objective for a translational CMV vaccine as it fails to evaluate cross-strain virus protection. A novel clinical strain of GPCMV (TAMYC) from the salivary gland of a commercial animal was successfully isolated [37]. This clinical strain has the capacity to reinfect animals convalescent for 22122 strain [24,37] and vaccines with success against 22122 have limited efficacy against TAMYC strain making it ideal for cross protection vaccine studies [17,23,27,38–40]. A significant problem with clinical CMV strains is that they are highly cell-associated and therefore capable of evading neutralizing antibodies that target cell free virus [1]. In HCMV, a T cell response is considered effective at controlling the levels of virus infected cells [41]. Convalescent HCMV patients produce a significant T cell response against viral tegument protein pp65 [42,43]. GPCMV encodes a functional pp65 homolog (GP83) involved in innate immune evasion targeting IFI16 cGAS-STING pathway. Although not necessary for direct entry, pp65 is an enhancement factor for endocytic cell infection and pathogenicity in animals [38,39,44]. GP83 is a T cell antigen in vaccinated or GPCMV infected animals [17,39,45].

GPCMV gB and pp65 proteins are strongly conserved between strains (99% and 100%) respectively [27,39]. In this report, we investigated the hypothesis that GPCMV cross-strain vaccine efficacy could be improved by combining trimeric capable gB (antibody target) and pp65 (T cell antigen) in a recombinant Ad vaccine strategy (AdgB+AdGP83). Novel GPCMV gB antibody ELISAs against prefusion gB and a homolog AD-1 gB domain were included to more comprehensively define the GPCMV gB immune response. Additionally, gB antibody ELISAs to the AdgB+AdGP83 vaccine was compared to historical sera from a prior AdgB vaccine [23] as well as GPCMV (22122) natural convalescent immune sera. Neutralizing antibody titers were evaluated for both 22122 and TAMYC strains. Concurrent neutralization assays were compared to historical AdgB vaccine sera and GPCMV convalescent sera with results indicating that the combined vaccine had a more limited GPCMV neutralizing titer likely associated with lower gB (AD-1) titer. Combined vaccine strategy induced a specific cell mediated response to GP83 in a splenocyte IFNγ ELISPOT assay. In a horizontal vaccine protection study, animals in the combined vaccine group exhibited protection against challenge virus (clinical strain TAMYC) compared to unvaccinated control animals with reduced viral loads in tissues but limited impact on viremia or salivary gland infection. Overall, results indicate that a combined AdgB+AdGP83 vaccine poorly improves cross-strain protection compared to a standalone trimeric gB vaccine. Potentially, reduced pp65 expression by TAMYC strain compared to 22122 prototype strain enables T cell evasion. Additionally, pp65 as an IFI16/ cGAS-STING innate immune evasion factor negatively impacts gB vaccine antibody response in an undefined mechanism compared to standalone gB vaccine as suggested by a reduced response to immunodominant gB(AD-1) domain. Results indicate that other viral T cell antigens should preferentially be used in conjunction with gB in a combined CMV vaccine strategy for enhancement of protective immunity.

## 4. Materials and Methods

### Cells, viruses, oligonucleotides and genes

GPCMV was propagated on various cell lines. Guinea pig fibroblast lung cells (GPL; ATCC CCL 158) and renal epithelial cells (REPI) for specific tropism studies as previously described [16,46]. Clinical GPCMV strain TAMYC [37] and prototype 22122 strain (ATCC VR682) were expanded from separate guinea pig salivary gland stocks and exclusively propagated on REPI cells for maintenance of fully tropic virus stocks [16]. Recombinant defective adenovirus (Ad5) vectors (E1 and E3 deleted) were previously described for gB and pp65 [23,39]. ORFs were under HCMV IE enhancer control with 3’ SV40 polyA sequence. High titer CsCl gradient purified recombinant defective adenovirus virus stocks (10^12^ TDU/ml) were generated by Welgen Inc. (MA). All oligonucleotides were synthesized by Sigma-Genosys (The Woodlands, TX).

### Ethics

All guinea pig animal studies were carried out under IACUC permit (Texas A&M University). All study procedures were carried out in strict accordance with the recommendations in the Guide for the Care and Use of Laboratory Animals of the National Institutes of Health. Studies were carried out in female guinea pigs. Animals were observed daily by trained animal care staff, and animals that required care were referred to the attending veterinarian for immediate care or euthanasia. Terminal euthanasia was carried out by lethal CO_2_ overdose, followed by cervical dislocation, in accordance with IACUC protocol and NIH guidelines. Animals purchased from Charles River Laboratories were verified as seronegative for GPCMV by toenail clip bleed and anti-GPCMV ELISA of sera, as previously described [39]. Animal studies were carried out to determine: (i) the immune response to the AdgB+AdGP83 vaccine; (ii) the level of protection provided by AdgB+AdGP83 vaccine against wild-type GPCMV challenge (TAMYC strain) compared to the no-vaccine control. Historical sera used in comparative studies for ELISAs and neutralization assays were obtained from previous studies: GPCMV (22122 strain) convalescent animal sera [24]; AdgB vaccinated animal sera [23].

### ELISAs

Anti-GPCMV ELISA was carried out as previously described [15] to determine GPCMV sero-status of commercially obtained guinea pigs (Charles River) as commercial animal colonies are not all negative for GPCMV. Specific glycoprotein gB ELISAs were performed as previously described [15,17]. Novel gB glycoprotein antibody ELISAs were included for: (1) prefusion gB (prefgB) ; and (2) gB(AD-1) domain. Based on the alignment of GPCMV gB with HCMV gB protein amino acid sequences, a GPCMV prefgB was generated by gene synthesis (Genscript) incorporating into the ORF removal of the furin site and additional specific amino acid substitutions equivalent to those made in the HCMV gB ectodomain to generate a locked version of GPCMV prefgB [24,47]. The ORF also incorporated a C-terminal FLAG epitope tag. The GPCMV prefgB was cloned into the pcDNA3.1(+) vector (Invitrogen) under HCMV IE enhancer promoter control to enable transient expression in plasmid-transfected cells to generate a coating antigen and expression of prefgB was verified using Western blotting [24]. (2) A homologous gBAD-1 domain was previously identified between HCMV gB and various animal CMV gB proteins based on BLAST alignment [24,48]. The minimal gB(AD1) region was extended for GPCMV gB with additional flanking N- and C-terminal amino acids to match the extended AD1 sequence used to express a commercial HCMV gB(AD-1) protein as a GST-gB(AD-1) fusion protein (Sigma-Aldrich). A synthetic GST protein ORF encoding the C-terminal fused GPCMV gB(AD-1) region (gB amino acid codons 524-646) was cloned into a bacterial expression vector and expressed as a recombinant protein (Genscript). The recombinant protein was purified using GST affinity chromatography for use in gB(AD-1) ELISA studies. For gB(AD-1) ELISA, the purity of the recombinant gB(AD-1) protein was verified using SDS-PAGE, Coomassie gel staining, and Western blotting [24]. Both antigens were determined for optimal plate coating concentrations: A) 2 mg/ml prefgB; B) 3 ug/ml gB(AD-1). The net OD (absorbance at 450 nm) was obtained by subtracting the OD of Ag− from that of Ag+. ELISA reactivity was considered positive if the net OD was ≥0.2, as determined using GPCMV-negative serum. Assays on pooled sera were conducted a minimum of three times, and the mean titer was determined. In addition, varying coating concentrations of the control carrier recombinant GST protein (Genscript) were tested for background using both GPCMV seropositive and seronegative control sera [24].

### Neutralization assays

GPCMV neutralization assays (for determination of the NA_50_) were performed on GPL fibroblast and renal epithelial cells with PC+ GPCMV stocks generated on a matching cell line as previously described [17]. GPCMV NA_50_ assays were carried out separately on 22122 and TAMYC strains. Neutralization assays were carried out with pooled sera from a specific group. The final neutralizing antibody titer was the inverse of the highest dilution producing a 50% or greater reduction in plaques compared to that for the virus-only control. Assays were performed from each sample three times and performed concurrently with the same virus stocks between groups.

### IFNγ ELISPOT

The guinea pig IFN-γ enzyme-linked immunospot (ELISPOT) assays were performed by following a previously described protocol using freshly isolated splenocytes and GPCMV GP83 peptide pools [49]. Anti-guinea pig IFN-γ monoclonal antibodies (V-E4 and N-G3) were previously characterized [50]. Briefly, polyvinylidene difluoride (PVDF) membrane 96-well plates were coated with a guinea pig IFN-γ capture antibody, V-E4, incubated overnight at 4°C, and blocked, and then freshly isolated splenocytes were added before being exposed to GP83 GPCMV 9mer peptide pools spanning the entire ORF and incubated for 18 h. A biotinylated detection antibody, N-G3, was added before streptavidin-AP conjugate (R&D Systems) and then detected with BCIP/NBT (5-bromo-4-chloro-3-indolylphosphate–nitroblue tetrazolium) (Life Technologies). Membranes were dried before spots were counted on an ImmunoSpot S6 system (CTL). Final counts were calculated based on the number of spot-forming cells (SFC) per 10^6^ cells after background spots (cells only without any stimulation) were subtracted. Concanavalin A (ConA; 10 μg/ml) was used as a positive control, and other controls included cells-only control, dimethyl sulfoxide (DMSO) control (peptide background), and nonspecific peptide control and medium-only control.

### Western blot assays

Western blots were carried out on cell lysates separated by 4-20% SDS-PAGE under denaturing conditions and performed as previously described [15,16,23]. Primary antibodies were GPCMV-specific antibodies (rabbit or mouse derived). The GPCMV gB antibody (mouse monoclonal) was a gift (Dr. Britt, UAB) [51]. Custom rabbit antibodies to pp65 (GP83) and pp71 (GP82) were previously described [39]. Anti-epitope tag antibodies included FLAG (Novus Biological) and His (Bethyl) [16,17]. Primary antibodies were used at 1/1000 dilution. Secondary antibodies: anti-mouse or anti-rabbit IgG HRP-linked secondary antibodies for western blot (Cell Signaling) were used at 1/2000 [17].

### qPCR

Tissues were collected from the euthanized guinea pigs to determine the viral load. For tissue DNA extraction, FastPrep 24 (MP Biomedicals) was used to homogenize the tissues as a 10% weight/volume homogenate in Lysing Matrix D (MP Biomedicals). DNA was extracted using the QIAcube HT (Qiagen) according to the manufacturer’s tissue protocol instructions. Viral load was determined by real-time PCR on a LightCycler 480 (Roche Applied Science), as previously described [16,49]. Primers and hydrolysis probes were designed using the LightCycler Probe Design2 program to amplify a product from the GPCMV viral polymerase subunit GP44 gene: forward primer 5’TCTCCACGGTGAAAGAGTTGT; Reverse primer 5’GTGCTGTCGGACCACGATA; hydrolysis probe 5’FAM-TCTTGCTCTGCAGGTGGACGA-BHQ1. Data were analyzed using the LightCycler Data Analysis Software (version 1.5.1; Roche). A standard curve was generated using serial dilutions of the GPCMV 22122 GP44 plasmid at known concentrations for quantification and assay sensitivity. The same standard curve was generated using TAMYC GP44 plasmids to demonstrate that the Universal GP44 primer-probe set could detect both strains. The sensitivity of the assay was determined to be two copies per reaction. Viral load was expressed as copy number/mg tissue or copies/ml of blood. The results were calculated as the mean value of all positive results. Positive triplicates were considered positive. Assays were repeated for those with any wells below the level of detection. Results were considered positive if at least 2/3 wells were positive on repeated runs. Wells with extrapolated data below the level of detection were not included in the mean calculation as positive results.

### Statistical analysis

All statistical analyses were conducted with GraphPad Prism (version 7) software. Fisher’s exact test, Student’s t test (unpaired) with significance taken as a *p* value of <0.05 or as specified in the figure legends.

## 5. Results

### Evaluation of immune response

The ability of combined Ad based vaccine to trimeric capable gB and pp65 antigens (AdgB+AdGP83) were evaluated for cross-strain protection against clinical strain TAMYC challenge in a non-pregnant pathogenicity model. An outline of the study is shown in Figure 1. Seronegative dams (n=12) at days 0, 28 and 56 were vaccinated SQ (vaccine dosage 10^8^ Transduction Units (TDU)/shot per Ad vector) with bleeds at days 26, 54 and 78 for evaluation of anti-gB(wt) antibody response (Figure 1A) [23]. Verification of gB (AdgB) and pp65 (AdGP83) expression by recombinant Ad vectors was confirmed by Western blot of cell lysate of fibroblast cells transduced with specific Ad vector (Figure 1B and C). Anti-gB ELISA results demonstrated that all vaccinated animals had a response to gB antigen and that average titer increased between first, second and third vaccination (Figure 1D). At day 78 post initial vaccination, antibody immune response was further characterized with a series of additional gB ELISAs: (1) anti-gB(wt); (2) prefusion gB; and (3) gB (AD-1 domain). Results from vaccinated (AdgB+AdGP83) pooled sera were compared with historical pooled convalescent sera from single shot 22122 infected animals [24] that mimicked natural infection response (see Figure 2). Additionally, combined vaccine response was also compared to historical pooled sera from AdgB only vaccinated animals [23] (see Figure 2).

**Figure 1.**
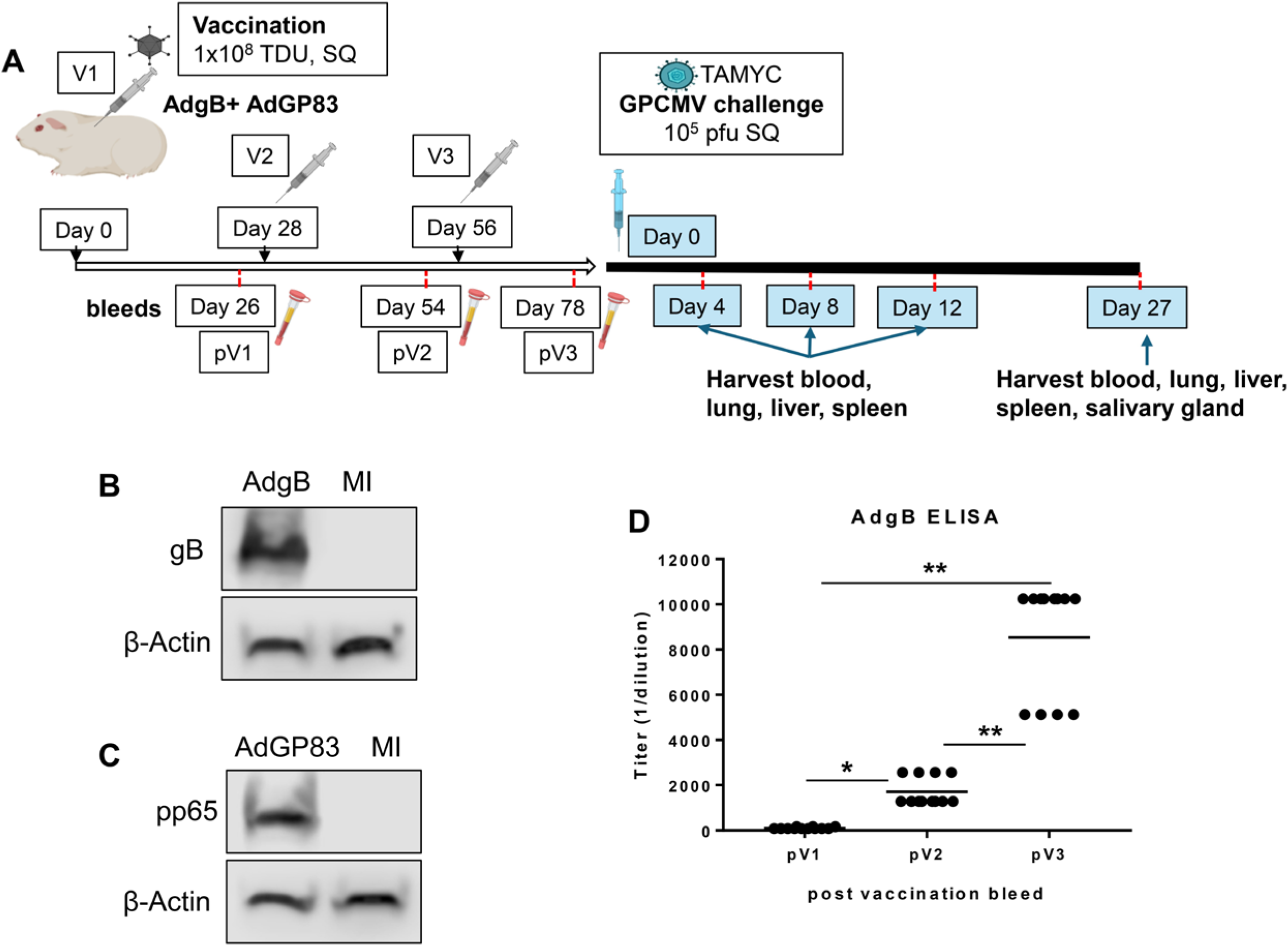
Schematic overview of vaccine protection study against GPCMV TAMYC strain challenge. (A) Seronegative animals (n = 12) were vaccinated 3 times (10^8^ PFU) via SQ route on days 0, 28, and 56. Serum was collected 3 to 4 weeks after each vaccination (days 26, 54, and 78). At day 100, animals were challenged with TAMYC strains (10^5^ pfu, SQ route) then viral load in the target organs or blood evaluated by real-time PCR on days 4, 8, 12 and 27 post challenge. (B-C) Guinea pig fibroblast cells were transduced with AdgB (B) or AdGP83 (C) to confirm antigen expression by Western blot analysis of cell lysate with gB or pp65 specific antibodies. (D) Sera collected after each vaccination round (pV1, pV2, pV3) were assayed for gB titer by gB-specific ELISA. Graph plotted for each individual animal at different post-vaccination time points with mean value represented by solid black line. Statistical analysis was determined by unpaired Student’s t test, * *p* < 0.05; ** *p* < 0.0001.

**Figure 2.**
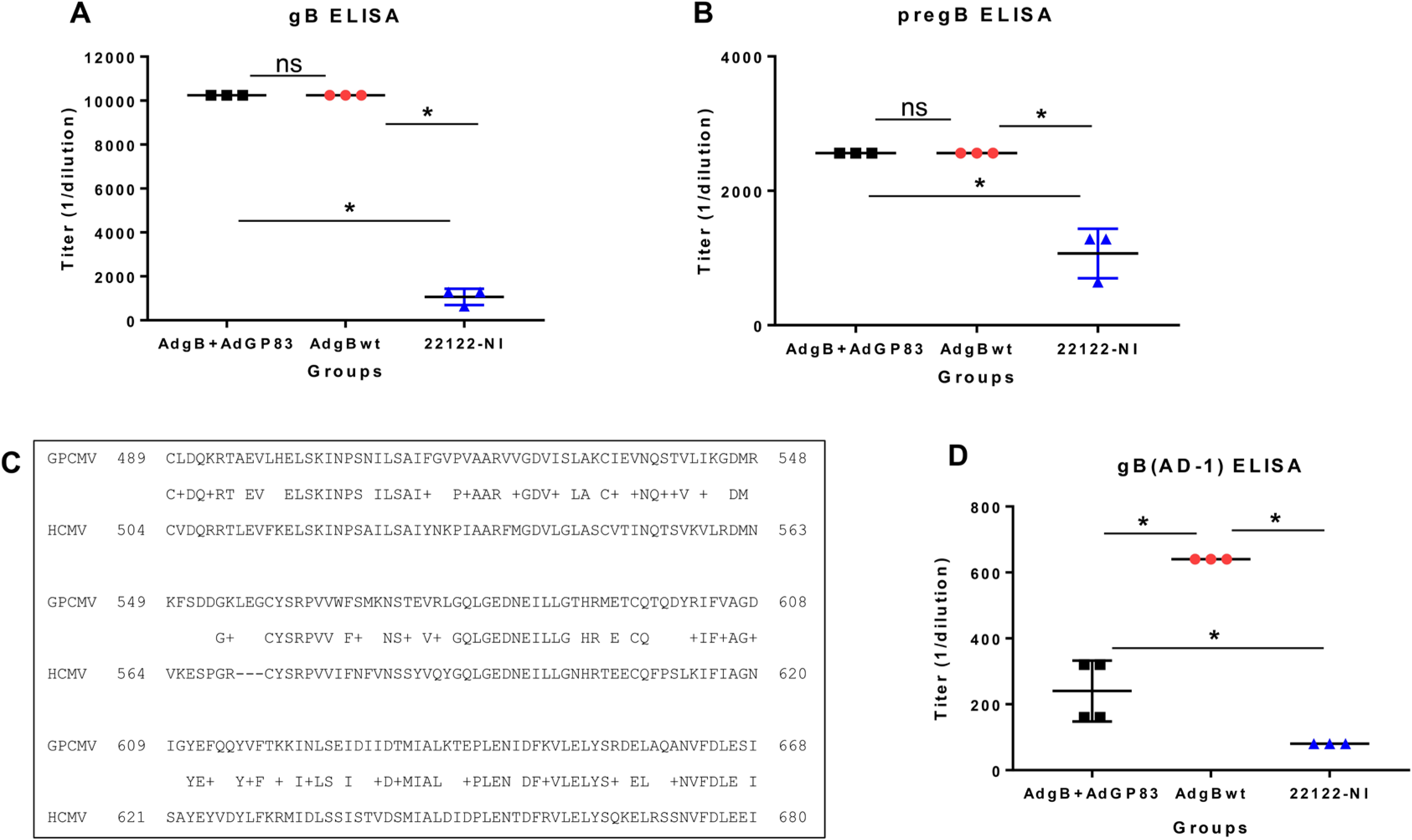
Antibody immune responses to AdgB+AdGP83 vaccine compared to historical vaccine sera (AdgB) or GPCMV convalescent sera. Pooled animal sera of AdgB+AdGP83 (black) was evaluated for antibody response to gB with specific ELISAs with comparisons to concurrent assays of historical sera from gB only vaccine AdgB (red) [23] or 22122 GPCMV convalescent sera (blue) [24] for antibody titers to specific target ELISAs: (A) anti-gB or (B) anti-prefusion gB. (C) BLAST amino acid alignment of the predicted GPCMV gBAD1 domain with HCMV gBAD1 [24]. (D) anti-gB(AD1) ELISA of pooled sera collected from AdgB+AdGP83 (black), AdgB (red), or 22122 GPCMV convalescent sera (blue). All assays were repeated minimum of 3 times. Statistical analysis was determined by unpaired Student’s t test, \**p* < 0.05; ns= not significant.

The anti-gB(wt) ELISA titer response (trimeric capable gB) in the gB+pp65 vaccine group was similar to the titer of historical AdgB only sera (Figure 2A) with anti-gB titers ∼nine-fold higher than that of GPCMV(22122) convalescent sera (mean titer: 10240 vs 1067) which was statistically significant (Figure 2A). Prefusion gB is a newly defined antigen for HCMV [47] and recently identified for GPCMV [24]. A synthetic codon optimized gene encoding prefgB was generated (Genscript) and cloned into a mammalian expression plasmid used to generate ELISA coating prefgB antigen as previously described [24]. Comparison of anti-prefusion gB ELISA results between groups with pooled sera demonstrated that both the combined gB+pp65 vaccine and gB only vaccine group had similar higher titer immune response to prefgB compared to that of convalescent (22122) immune response (Figure 2B). The gB(AD-1) domain is considered the immunodominant region in HCMV gB and a homolog domain was recently characterized for GPCMV gB [24]. Figure 2C shows BLAST alignment of the predicted GPCMV gB(AD1) domain with HCMV gB(AD1). Recombinant GPCMV gB(AD-1) fused to the C-terminus of the GST carrier protein was expressed in bacteria and purified for use as a coating antigen as recently described [24]. A control ELISA with recombinant GST protein (Genscript) lacked activity with GPCMV seropositive and seronegative sera, demonstrating a lack of pre-existing antibodies to GST protein in animals (data not shown) [24]. A comparison anti-gB(AD-1) ELISA titers between groups indicates that the highest immune response was obtained for gB only vaccine with ∼3-fold higher titer than combined AdgB+AdGP83 vaccine strategy (Figure 2D). The lowest antibody titer to gB(AD-1) was obtained in the GPCMV(22122) convalescent group (mean titer 80) and was statistically lower than titers obtained for other groups. Overall, results indicated that although the combined AdgB+AdGP83 vaccine strategy had similar anti-gB(wt) and prefgB ELISA titers, it had drastically reduced the immune response to gB(AD-1) domain with a titer approximately double that of convalescent sera.

Next, sera from combined vaccinated animal group were evaluated for neutralizing antibody titers (NA_50_) against 22122 and TAMYC GPCMV strains. Pooled sera from the combined vaccine group were concurrently evaluated for virus neutralization with historical pooled sera from AdgB vaccine group [23] or convalescent sera (22122 infected animals) [24]. Neutralization assays were carried out on both fibroblast (GPL) and epithelial (REPI) cells (Figure 3) using pooled sera collected at 78 days after the initial vaccination for gB+pp65 vaccine group. Evaluation of GPL NA_50_ for 22122 strain (Figure 3A) demonstrated significant differences between gB+pp65 vaccine group and prior AdgB with neutralization titers highest for gB only vaccine. The gB+pp65 vaccine sera also had a statistically lower NA_50_ titer than convalescent 22122 sera (Figure 3A). This general trend against 22122 strain was also observed for NA_50_ on epithelial cells but the difference between titers for gB+pp65 vaccine sera and convalescent sera was not significant. Historical gB only vaccine sera also exhibited higher NA_50_ titers against TAMYC strain compared to AdgB+AdGP83 vaccine sera on both fibroblast and epithelial cells (Figure 3C and 3D) with overall lower neutralizing titers. TAMYC strain NA_50_ titers were not statistically different between convalescent sera (22122) and AdgB+AdGP83 vaccine sera on both fibroblast and epithelial cell types. Overall, the NA_50_ results indicated that the historical gB only vaccine sera had better neutralizing capability against both strains of GPCMV on both fibroblast and non-fibroblast cell types compared to combined AdgB+AdGP83 vaccine strategy and convalescent (22122) sera.

**Figure 3.**
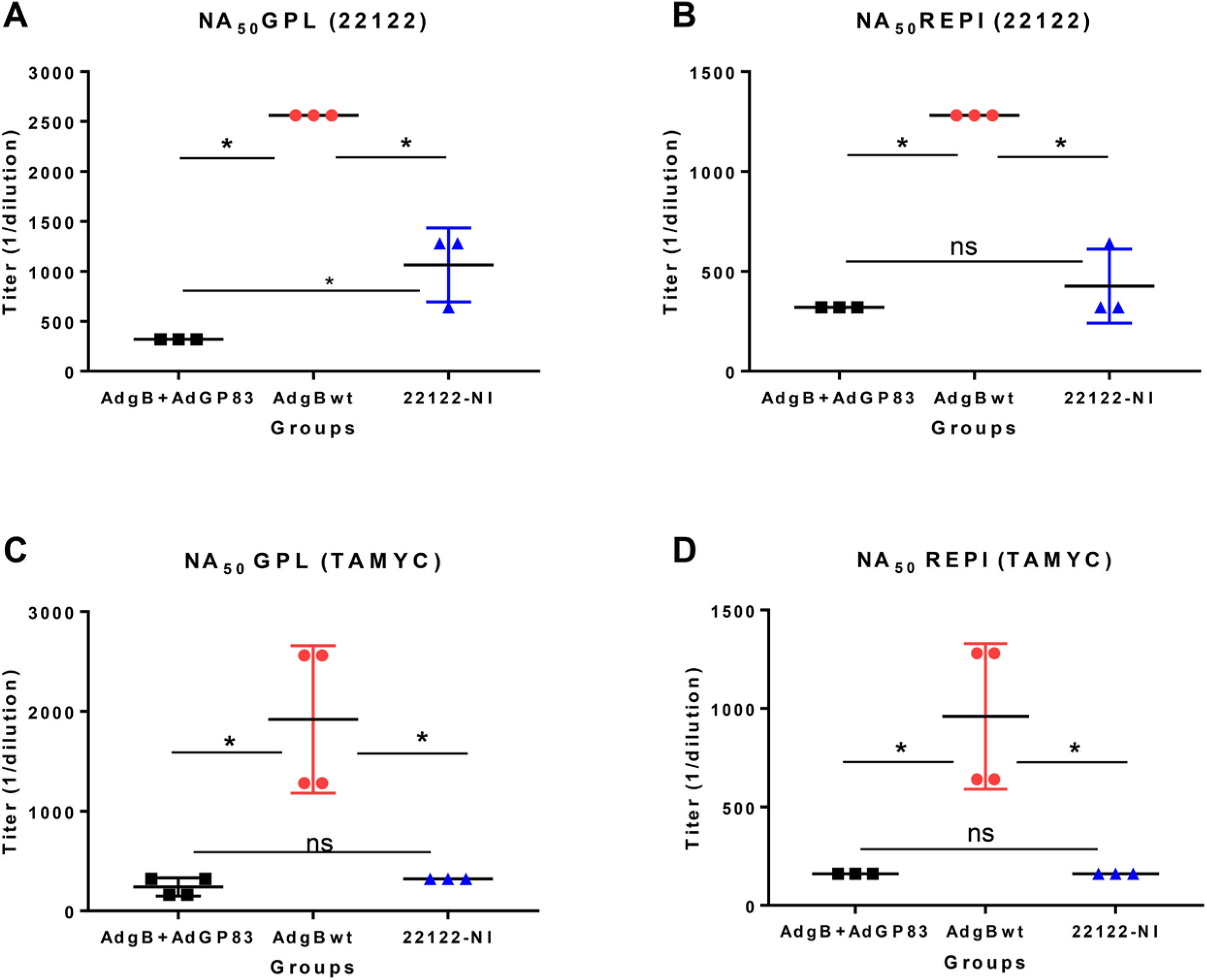
GPCMV neutralization by anti-AdgB+AdGP83 compared to historical sera from anti-AdgB or GPCMV (22122) natural convalescent sera (22122-NI). Pooled animal sera of AdgB+AdGP83 (black) or AdgB (red) vaccinated animals were compared to pooled sera of GPCMV (22122) convalescent sera (blue) animals for neutralization of 22122 or TAMYC strain GPCMV on different cell lines: (A) Neutralization of 22122 on GPL fibroblasts cells; (B) Neutralization of 22122 on epithelial cells (REPI); (C) Neutralization of TAMYC virus on GPL cells; (D) Neutralization of TAMYC on epithelial cells (REPI). All assays were repeated minimum of 3 times. Statistical analysis was determined by unpaired Student’s t test, \**p* < 0.05; ns= not significant.

In addition to antibody response to gB, the combined AdgB+AdGP83 vaccine strategy was expected to induce a cell mediated response to the GPCMV pp65 tegument protein with the predicted amino acid sequence being 100% conserved between GPCMV strains [39]. IFNγ-ELISPOT assay of splenocytes using overlapping peptides corresponding to GP83 ORF demonstrated a positive cell mediated response when splenocytes were stimulated with specific GP83 peptide pools (Figure 4A) which was similar to that previously observed for a AdGP83 only vaccine [39]. Results in Figure 4B show ELISPOT images for GP83 peptide stimulated splenocytes as well as negative peptide pools. Results also show splenocyte ELISPOT results for positive control ConA stimulated cells as well as unstimulated cells (Figure 4B). Positive stimulating GP83 peptide 9mer sequences are boxed in Figure 5 and demonstrated that there were 18 regions spread across the 365 amino acid pp65 protein sequence that stimulated a response. However, there was an extended region within pp65 ORF that induced a positive response between amino acids 337-389. Overall, results indicated that the combined vaccine strategy induced a cell mediated response to the pp65 antigen.

**Figure 4.**
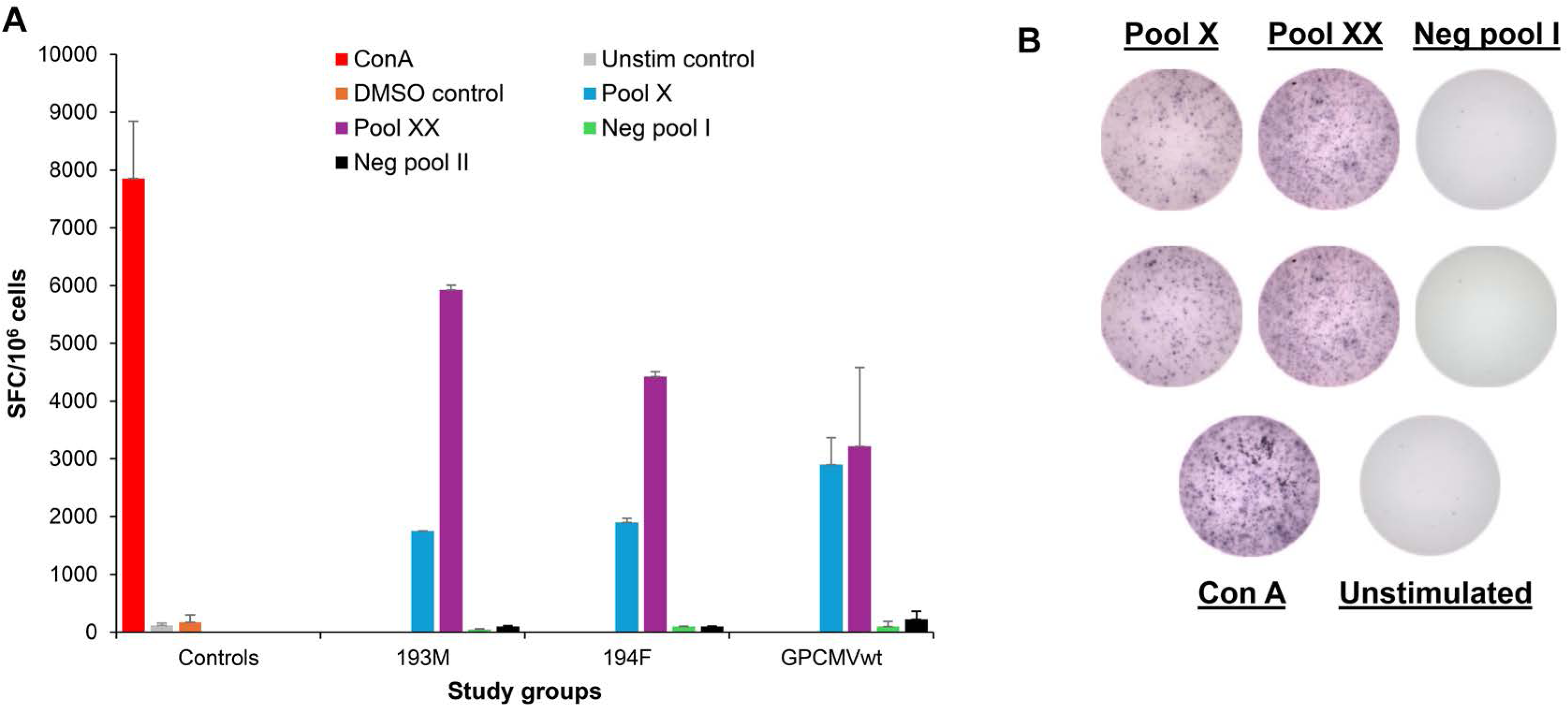
GP83 (pp65) T cell response to the AdgB+AdGP83 vaccine determined by IFN-ELISPOT assay. (A) Guinea pig-specific IFNγ response to GPCMV GP83 peptide pools in AdgB+AdGP83 vaccinated animals or wtGPCMV^+^ convalescent animals are shown. Two reactive GP83 peptide pools, peptide pools X (blue) and XX (purple), previously identified [39], responded with splenocytes isolated from AdgB+AdGP83 vaccinated animals or seropositive wtGPCMV-infected animals. Red, ConA positive control; orange, DMSO control; gray, unstimulated (unstim) control; green, unresponsive negative (neg) peptide pool I; black, unresponsive negative peptide pool II. Final counts were calculated based on the number of SFC per 10^6^ cells after the values for background spots (cells only, without any stimulation) were subtracted. (B) Selected representation of ELISPOT positive (pool X and pool XX) and negative (neg pool I) well readouts from IFNγ assay. ConA positive control and unstimulated wells are also included.

**Figure 5.**
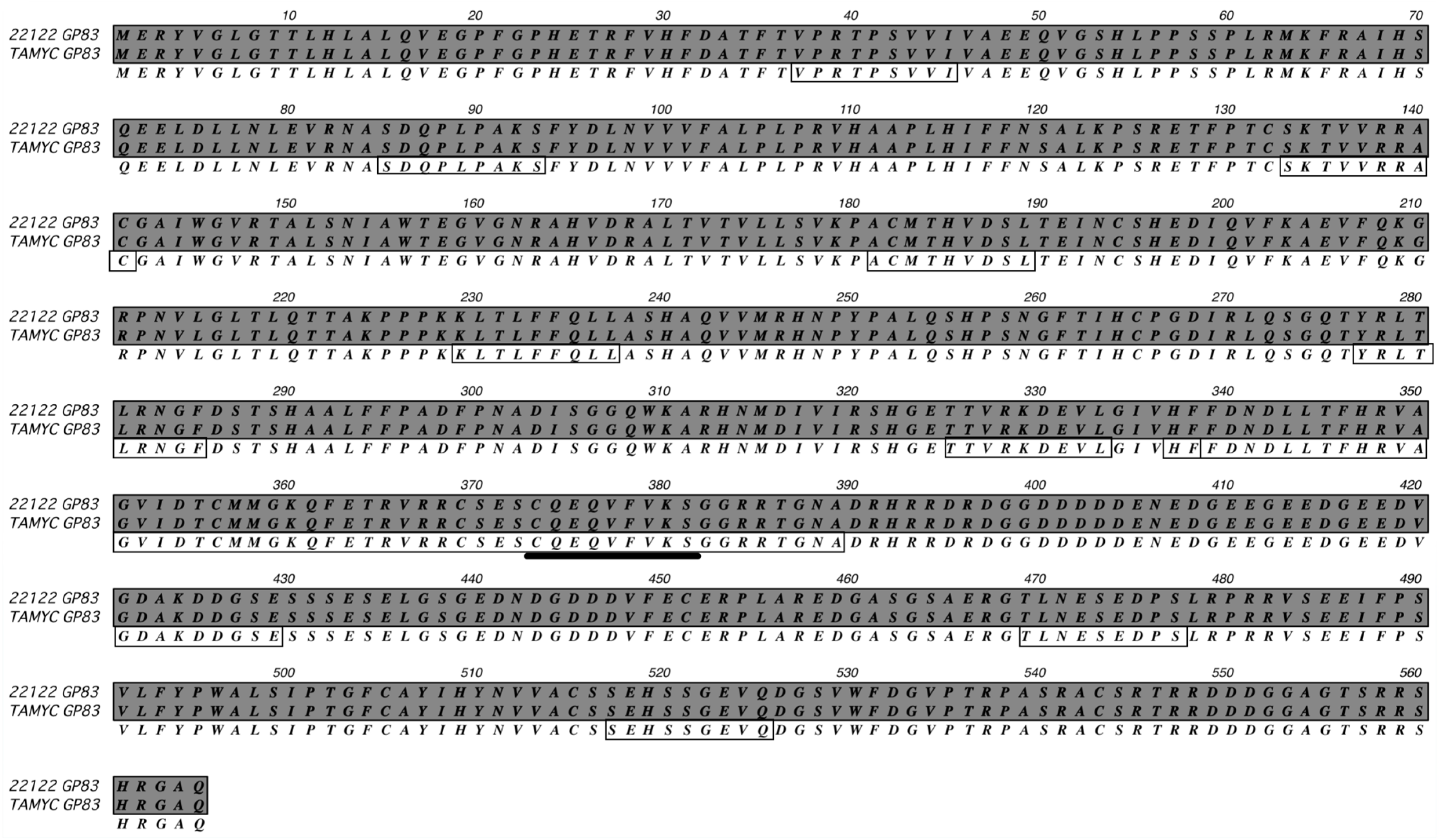
Comparative alignment of the predicted GP83 (pp65) protein amino acid sequences from GPCMV strains 22122 and TAMYC. ClustalW alignment of the predicted amino acid sequences of GP83 from GPCMV strain TAMYC [39] and strain 22122 (GenBank accession no. AB592928) is shown. Gray background with consensus letter in the third row indicates amino acid sequence identity. Boxed sequences represent the peptides in the reactive pools X and XX with bold underlined representing overlapping sequence within the pools. Sequence alignment was carried out using DNAstar software.

### GPCMV gB+pp65 vaccine protection outcome against TAMYC strain challenge

In a GPCMV challenge pathogenicity protection study, AdgB+AdGP83 vaccinated animals (n=12) were challenged with GPCMV TAMYC strain (10^5^ pfu, SQ). A GPCMV seronegative group (group 2, n=12) was similarly challenged with TAMYC strain virus. At 4, 8, 12 and 27 days post-vaccination (DPC), animals in both groups were randomly euthanized (n=3/group) and viral load in target organs and blood evaluated (lung, liver, spleen and blood). Additionally, at 27 DPC, salivary gland tissue was also evaluated for viral load. Results (Figure 6) demonstrated that AdgB+AdGP83 vaccine was effective in reducing viral load in target organs (liver, lung, spleen) at all time points compared to the unvaccinated control group. Viral load in the salivary gland was reduced by over 2 logs at 27 DPC from >10^6^ genome copies/mg tissue in control to less than 10^4^ genomic copies in the vaccine group (Figure 4D). The AdgB+AdGP83 vaccine significantly reduced viremia to be seen at 4, 8 and 12 DPC (Figure 4E) compared to viremia in the unvaccinated control group. Comparison of the two study groups showed statistically significant differences (*p* < 0.05) in viral load in all tissues and blood at all time points. Overall, results indicate that the immune response in vaccinated animals was effective in limiting virus dissemination but crucially the vaccine did not prevent viremia or dissemination of virus to the salivary glands, which were considered significant shortfalls in vaccine efficacy. Potentially, TAMYC strain has an ability to partially evade T cell response to pp65 antigen by low level expression of the antigen in both fibroblast and epithelial cells compared to prototype strain 22122. Figure 7 demonstrates that in virus infected cells at 3 days post infection (MOI 1 pfu/cell) the level of GP83 (pp65) expression is >30 fold higher for 22122 compared to TAMYC strain and the same contrast is observed on both fibroblast and epithelial infected cells (Fig 7A and Fig 7B). In contrast, the lower tegument protein pp71 (GP82) [39] and gB are expressed at comparable levels between strains with similar levels on both fibroblast and epithelial cells for both viral strains (Fig 7A and Fig 7B).

**Figure 6.**
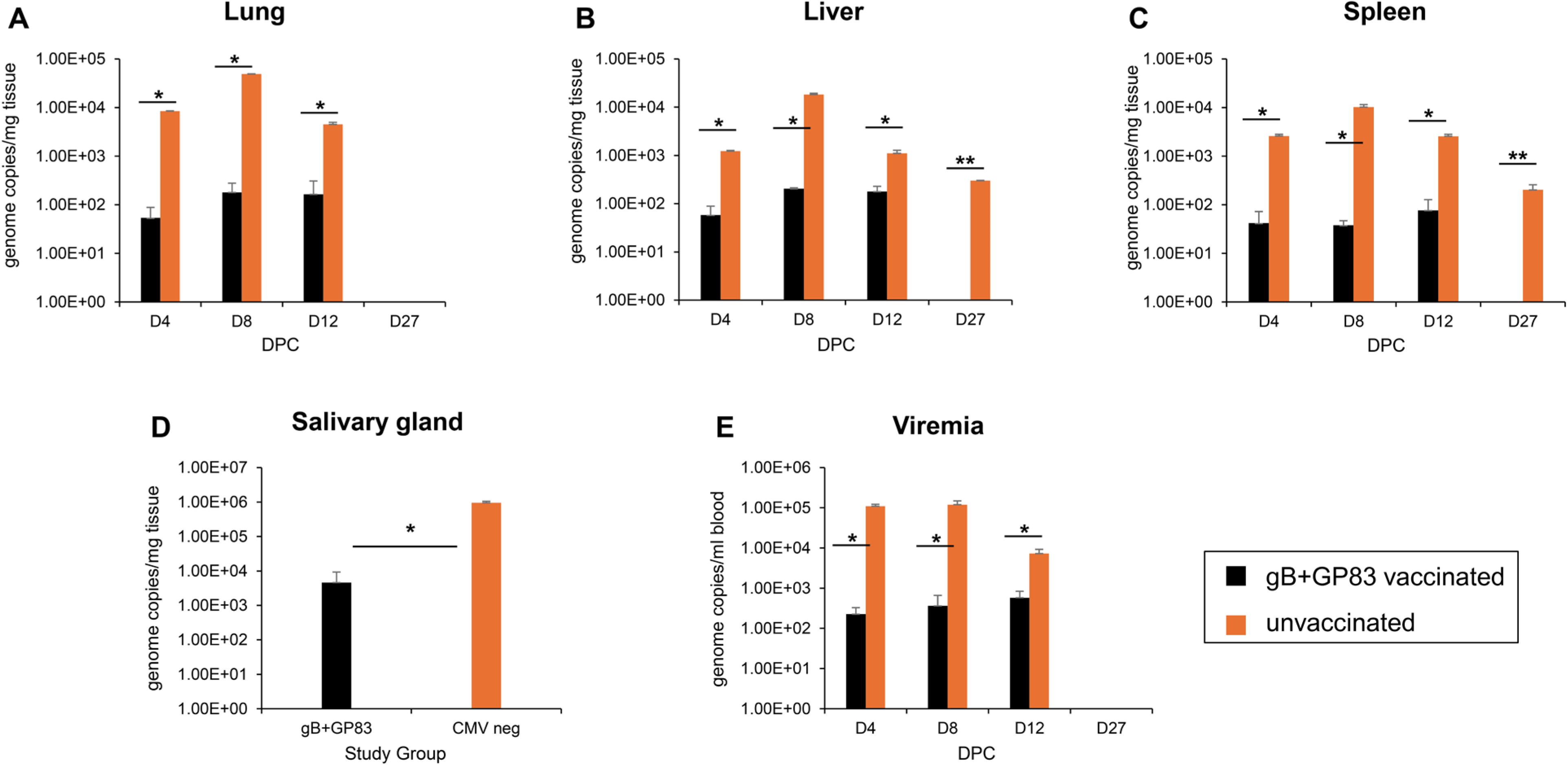
AdgB+AdGP83 vaccine protection against GPCMV (TAMYC strain) virus challenge. Pathogenicity study was conducted in female guinea pigs (*n* =12/group). Animals were vaccinated according to schedule in Figure 1A. AdgB+AdGP83 vaccinated group (black) or unvaccinated group (orange) challenged with TAMYC strain GPCMV (10^5^ pfu SQ). At 4, 8, 12, and 27 days post challenge (DPC), 3 animals per group were evaluated for viral load in target organs by real-time PCR of tissue extracted DNA. Viral load is plotted as the mean number of viral genome copies/mg tissue. Salivary gland tissue was evaluated only at day 27. (A) Lung; (B) liver; (C) spleen; (D) salivary gland (evaluated only at 27 DPC). (E) Viremia at 4, 8, 12, and 27 dpi is plotted as the mean number of genome copies/ml blood. Statistical analysis was performed by Student *t* -test. * *p <*0.05; ** viral load in vaccine group is below level of detection.

**Figure 7.**
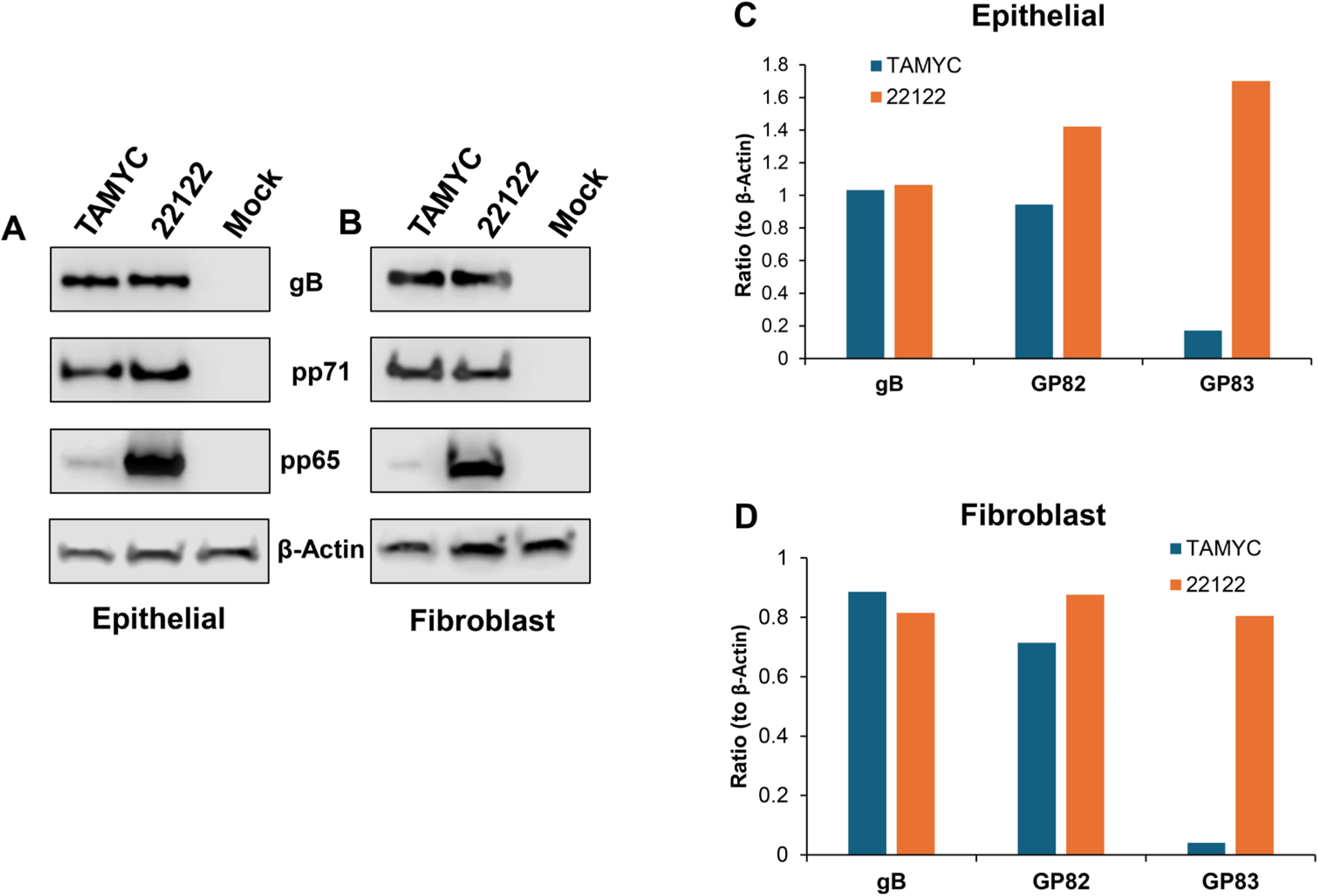
GPCMV gB, pp71 (GP82) and pp65 (GP83) expression in epithelial vs fibroblast cells. (A-B) Western blot analysis of gB, pp71 and pp65 expression by GPCMV strains (22122 vs TAMYC). Infection (MOI = 1pfu/cell) on different cell types: (A) epithelial cells or (B) fibroblast cells 2 days post infection. β-actin shown for loading control. Lane 1: GPCMV TAMYC strain infection; Lane 2: GPCMV 22122 strain infection; Lane 3: mock infection. (C-D) Each Western blot band was measured for intensity using the Image Studio (Licor) quantitative densitometry feature then normalized to β-actin. Comparison of gB, pp71, and p65 normalized expression in TAMYC (blue) vs 22122 (orange) infected epithelial (C) or fibroblast (D) cells.

## 6. Discussion

Despite more than 50 years of CMV vaccine research, a licensed vaccine remains an elusive goal as demonstrated by the recent failure of a Moderna mRNA-1647 CMV vaccine in phase III clinical trials [52]. Likely antibody and T-cell responses to key target antigens are required but the protective immune response of a CMV vaccine needs to greatly exceed that of natural CMV convalescent immunity as this immune response fails to prevent infection by a new viral strain. Consequently, a CMV vaccine needs to provide cross-strain protection since cCMV can occur as a result of both primary and non-primary infection [2]. Importantly, the level of cCMV related to non-primary infection is an underappreciated burden with increased cCMV rates occurring in countries or regions highly endemic for CMV [8,53]. The guinea pig represents the only small animal cCMV model and enables high through-put evaluation of vaccine candidates by evaluation of homolog target antigens [10]. The non-pregnant pathogenicity infection model studied in this report serves as a preliminary stage for evaluation of CMV candidate vaccines or intervention strategies prior to cCMV studies. Indeed, reducing or preventing virus dissemination in the mother is likely fundamental for protection against cCMV [54]. The majority of GPCMV vaccine studies have focused on protection against prototype strain 22122 in horizontal and vertical models. Although this is an important benchmark for vaccine success, the majority of prior studies fail to evaluate cross-strain virus protection which is of significant translational importance. A novel clinical strain of GPCMV (TAMYC) has the capacity to reinfect animals convalescent for 22122 strain. TAMYC strain is highly cell-associated similar to HCMV clinical strains [24,37]. Importantly, vaccines with prior success against 22122 virus have limited efficacy against TAMYC strain [17,23,27,38–40]. Although a trimeric gB vaccine was highly effective against 22122 strain [23], it lacked cross-strain protection against TAMYC strain despite 99% similarity in predicted gB amino acid sequence between strains [23,39]. The immunological basis for the limitation of GPCMV gB vaccine strategies in the guinea pig model are poorly defined in part because of limited characterization of gB antibody response and virus neutralization studies performed only on non-fibroblast cells. HCMV gB vaccine studies have identified specific antibody responses to domains within gB (AD-1 to AD-6) [55]. Furthermore, in HCMV, a prefusion confirmation of gB has recently been described but it is unclear if a specific response to this novel gB antigen enhances protection against the virus [47]. In the case of SARS-CoV2 and RSV, antibody response to prefusion glycoprotein antigens are the basis for vaccine efficacy [56,57].

In this report, additional aspects of the GPCMV gB antibody response were characterized and compared to natural convalescent GPCMV immunity and prior historical AdgB vaccine sera [23]. Novel gB antibody ELISAs included GPCMV gB(AD-1) domain and GPCMV prefusion gB [23]. In HCMV, the gB(AD-1) domain is necessary for oligomerization of gB [48] and presumably the case for GPCMV. HCMV gB(AD-1) domain antibody response is common in all convalescent patients and AD-1 is also a major target for neutralizing antibodies [32,58,59]. It is interesting to note that in HCMV clinical trials for both gB/MF59 and Moderna mRNA-1647 vaccines, gB(AD-1) binding antibodies were not detected [60,61]. Potentially, this impaired response contributed to limited Moderna vaccine efficacy in phase III clinical trials. Evaluation of historical sera from prior GPCMV AdgB vaccine study [23] demonstrated a robust antibody response to gB(AD-1) compared to sera from GPCMV convalescent animals. Sera from the combined gB+pp65 vaccine strategy despite inducing an anti-gB titer equivalent to prior AdgB vaccine sera had relatively limited response to AD-1 and was more similar to GPCMV convalescent sera (Fig 2). However, the combined vaccine strategy had an antibody response to prefusion gB more similar to AdgB only vaccine (Fig 2). The significance of a response to prefusion gB remains undefined for HCMV but viral prefusion glycoprotein antigens have been demonstrated to be important protective antigens eg. VZV gE [62]. Although the combined gB+pp65 vaccine strategy had a higher anti-gB titer than natural convalescent immunity, it is assumed that the majority of the response was directed towards undefined non-neutralizing gB antigens because the combined vaccine had lower neutralizing antibody titers to both 22122 and TAMYC strains on both fibroblast and epithelial cell lines compared to prior AdgB vaccine [23]. Previously, monomeric gB expressed in an Ad vector (AdgBTMD) produced anti-gB titers similar to trimeric capable gB (AdgB) but similarly had more limited neutralizing titers [23]. The target sites for non-neutralizing antibodies awaits further evaluation for GPCMV gB. In addition to neutralizing antibody activity, HCMV gB vaccines have ADCC and ADCP effects that enhance vaccine efficacy by targeting virus infected cells [61,63]. Unfortunately, a limitation of the current study is the lack of available guinea pig ADCC or ADCP assays that enables evaluation of this aspect of the GPCMV gB immune response, but this is worthy of future evaluation. However, similar to HCMV, GPCMV encodes viral receptors that bind antibodies that potentially enables evasion of ADCC response [64,65].

A CMV T cell response is important and more effective at controlling tissue specific viral load and virus infected cells compared to antibody response [41,66,67]. HCMV pp65 tegument protein is considered a major CMV T cell antigen and exploration of pp65 (GP83) vaccine strategies in the guinea pig model had varying levels of success against cCMV (22122 strain) with demonstrated IFNγ cell-mediated response in ELISPOT assays [17,45,68,69]. In a prior cCMV vaccine study, a combined monomeric gB plus pp65 vaccine strategy using recombinant LCMV vectors compared to gB only vaccine resulted in both vaccines attaining similar anti-gB titers [70]. However, this previous study provided limited insight into the gB antibody response and virus neutralization titers only evaluated on fibroblasts for 2212 strain [70]. Furthermore, the combined LCMV gB/pp65 vaccine approach did not improve protection against cCMV transmission (22122 strain) compared to gB only vaccine [70] and this was similarly the case for a MVA vector (gB monomer/pp65) vaccine approach [69]. An additional GPCMV study against cCMV with recombinant LCMV encoding pp65 or monomeric gB demonstrated that separately both antibody (gB) and T cell (pp65) antigen were similarly effective in reducing cCMV transmission (22122) compared to control unvaccinated animals [68] demonstrating the potency of both antibody and T cell response against 22122 strain. Consequently, it was surprising that limited protection against TAMYC strain virus was achieved against systemic infection given the high level of conservation between gB and pp65 proteins between strains and development of antibody and T cell response to respective viral antigens. Potentially, the significantly reduced level of pp65 antigen expression by TAMYC strain compared to 22122 strain (Figure 7) enables TAMYC virus to evade, in part, the T cell response. Additionally, the highly cell-associated nature of TAMYC strain results in lower level of cell-free virus compared to 22122 strain [37] and enables evasion of neutralizing antibodies.

It could be argued that the identified impaired antibody response to gB(AD-1) in the combined vaccine gB+pp65 vaccine strategy in this report has limited significance since a prior AdgB vaccine with enhanced prefgB and gB(AD-1) response compared to natural convalescent immunity lacked cross-strain protection against TAMYC virus [27]. But a trimeric capable AdgB vaccine strategy does exhibit cross-strain protection against cCMV [24]. Importantly, this current study demonstrates that a modified antibody response to gB occurs by inclusion of the pp65 antigen in a combined vaccine strategy. Potentially, this is attributable to the functional role of pp65 protein in both HCMV or GPCMV as a IFI16/cGAS STING pathway evasion factor [39] modifying innate and possibly adaptive immune response. This problem could be circumvented potentially by the use of non-functional pp65 antigen lacking ability to bind cGAS or IFI16 to enable a more comprehensive gB response while retaining a pp65 T cell mediated immunity. Alternatively, another viral T cell antigen could be used in conjunction with gB in a combined vaccine strategy. In HCMV infection, most convalescent patients evoke a T cell response to immediate early protein 1 (IE1) [71] which has demonstrated protective efficacy as a HCMV vaccine antigen in transplant patients [72]. GPCMV encodes a functional IE1 homolog, which induces as T cell response [73]. We recently demonstrated that IE1 exhibited efficacy against both horizontal and vertical GPCMV infection [73] and that IE1 antigen provided relatively high cross-strain protection against cCMV [73]. Furthermore, a combined gB and IE1 vaccine against TAMYC in both horizontal and congenital infection demonstrates high protective efficacy [24].

In summary, this study demonstrates limited cross-strain protection of a combined gB+pp65 vaccine strategy against GPCMV in a horizontal infection model despite evoking antibody (gB) and T cell response (pp65). Results indicate that inclusion of pp65 antigen altered gB antibody response and demonstrates the importance in further characterization of GPCMV gB response to place it on par with HCMV gB studies, especially if gB antigen is considered a corner stone of any CMV vaccine. Potentially, results suggest that the innate immune evasion function of pp65 has an undefined capacity to modify the adaptive immune response to gB possibly by direct interaction with IFI16 or cGAS. Consequently, other viral T cell target antigens might be better placed to complement gB immune response in a combined vaccine approach. Recent GPCMV studies demonstrate that IE1 is an important T cell antigen against GPCMV with capacity to synergize with gB immune response for enhanced protection against cCMV and is perhaps of higher priority for future studies. However, T cell antigens need to be expressed by the virus to enable effective targeting of virus infected cells and reduced pp65 expression by clinical TAMYC strain compared to 22122 prototype GPCMV demonstrates that potential for divergence and possible virus evasion strategy.

## 7. Author statements

### 7.1 Author contributions

K. Yeon Choi: Conceptualization, Methodology, Investigation, Data curation, Formal analysis, Writing - original draft, Writing - review & editing. Yushu Qin: Methodology, Investigation, Data curation, Formal analysis, Writing – original draft, Writing - review & editing. Nadia S. El-Hamdi: Methodology, Investigation, Data curation, Formal analysis, Writing – original draft, Writing - review & editing. Alistair McGregor: Conceptualization, Methodology, Investigation, Data curation, Formal analysis, Writing - original draft, Writing - review & editing, Funding acquisition.

### 7.4 Acknowledgements

The authors would like to thank Austin Selman for his technical assistance in some tissue culture experiments. Hybridoma cell lines for the anti-IFN-γ monoclonal antibodies were a generous gift from Dr. Schäfer (Robert Koch Institute, Germany).

### 7.5 Authorship

All authors attest they meet the ICMJE criteria for authorship.

